# Vorinostat Rescues SQSTM1 Palmitoylation and Restores Dysfunctional Autophagy in Huntington Disease

**DOI:** 10.64898/2026.03.01.708853

**Authors:** Yasmeen Alshehabi, Fatima Abrar, Milena Rabu, Anthony Dang, Firyal Ramzan, Dale D. O. Martin

## Abstract

Protein mislocalization, dysfunctional autophagy, and protein aggregation are key features of Huntington disease (HD). Accordingly, a central focus of our work has been identifying and correcting the protein mislocalization that drives the underlying autophagic defects which exacerbates protein aggregation. We previously demonstrated that palmitoylation, or S-acylation, of the autophagy receptor sequestosome 1 (SQSTM1; p62) is significantly reduced in the brains of HD patients and the YAC128 HD mouse model, thereby providing a possible mechanism for the cargo-loading failure observed in HD. Here, we identify the FDA-approved small molecule Vorinostat (suberoylanilide hydroxamic acid, SAHA) as a modulator of this pathway and show that it rescues SQSTM1 palmitoylation and restores autophagic function in HD models. Importantly, we demonstrate that Vorinostat crosses the blood-brain-barrier and significantly increases SQSTM1 palmitoylation in the cortices of YAC128 mice using acyl-biotin exchange and click chemistry assays. We further show that Vorinostat enhances autophagic flux, as evidenced by significant changes in autophagy marker levels and a marked increase in the colocalization of huntingtin with SQSTM1 and lysosomes. Finally, we investigate the mechanism of Vorinostat and propose a dual mode of action involving inhibition of depalmitoylating enzymes and transcriptional regulation of key pathway components. Collectively, these findings underscore SQSTM1 palmitoylation as a promising therapeutic target and support Vorinostat as a strong therapeutic candidate in HD.

## 1. Introduction

Protein aggregation is a hallmark of several neurodegenerative diseases, including Huntington disease (HD), amyotrophic lateral sclerosis (ALS), Alzheimer disease (AD), and Parkinson disease (PD)^1^. While the toxicity of aggregates remains debated, it is well established that the intracellular pathway responsible for their clearance, known as autophagy, is often impaired in these conditions^1–3^. As a result, autophagy has emerged as an attractive therapeutic target for promoting the removal of protein aggregates. However, only stimulating autophagy is insufficient, as it places additional stress on a system that is already compromised. Therefore, effective therapeutic interventions likely require addressing the underlying defects in autophagy while simultaneously increasing autophagic activity. Importantly, mislocalization is increasingly recognized as an early pathogenic event that contributes to proteostasis failure^3–5^, making the restoration of protein localization a growing focus for therapeutic development.

Macroautophagy, henceforth referred to as autophagy, involves the loading of cargo, such as toxic or aggregated proteins and damaged organelles, into a double-membrane structure known as a phagophore, which seals to form an autophagosome^6^ (Figure 1B). This autophagosome is then transported and fused with a lysosome to form the ultimate structure known as an autophagolysosome where the cargo is degraded with hydrolases^6^. The cargo-loading step is crucial, wherein sequestosome 1 (SQSTM1; also known as p62) can bind ubiquitinated cargo destined for degradation and deliver it to lipidated microtubule-associated protein 1 light chain 3 (LC3-II) located on the inside of the phagophore membrane^6,7^. In HD, autophagy is dysfunctional at several steps (Figure 1C)^8^. Critically, there is a cargo-loading failure that leads to an accumulation of empty autophagosomes and protein aggregates^9,10^.

**Figure 1.**
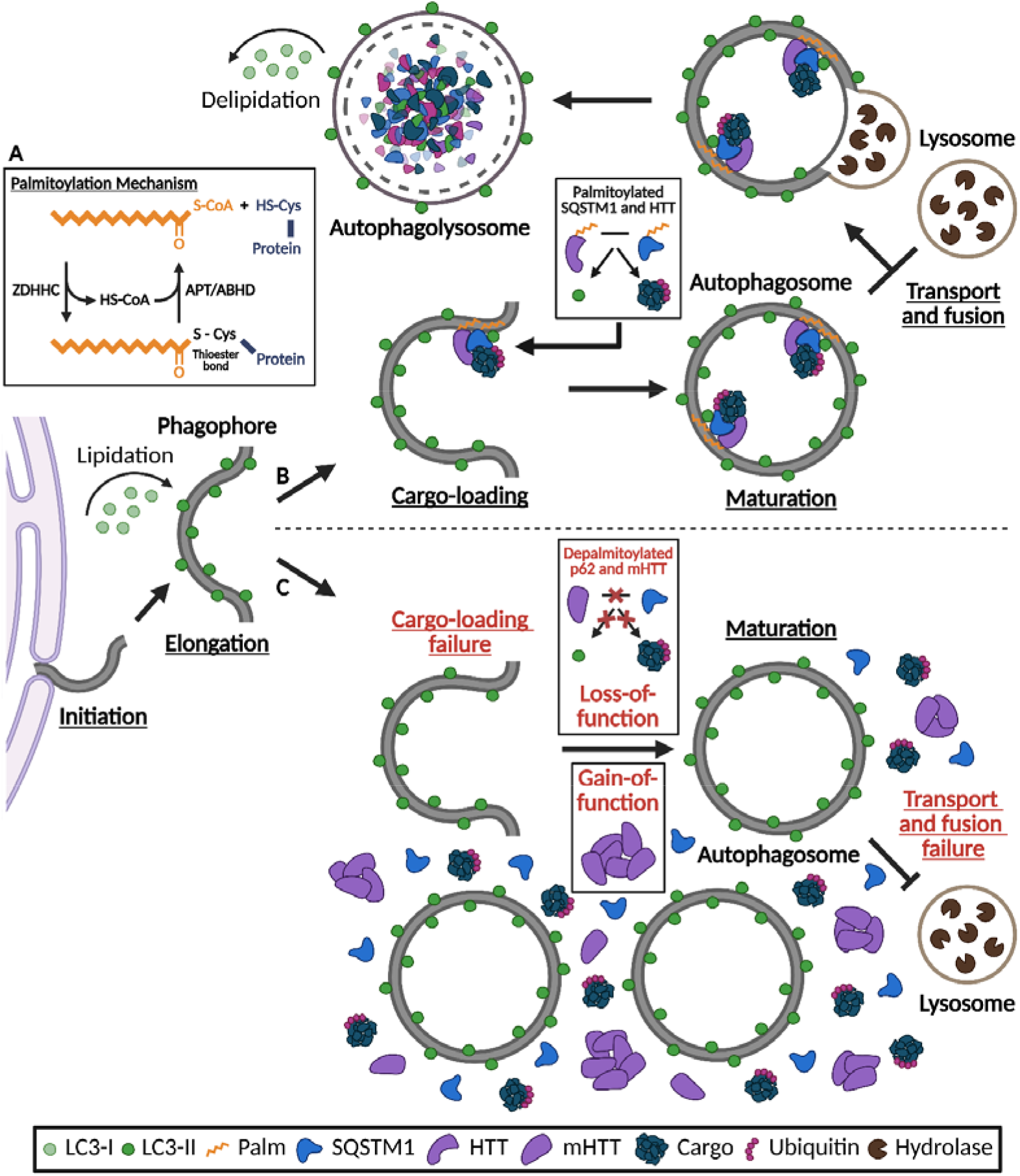
SQSTM1 and HTT palmitoylation may support healthy autophagy, and its deficiency may contribute to autophagic dysfunction in HD. **(A)** Palmitoylation involves the formation of a thioester bond between palmitoyl-CoA and a free cysteine residue in a protein via palmitoyl acyltransferases (ZDHHCs). Depalmitoylation involves the cleavage of a thioester bond via acyl protein thioesterases (APTs) or α/β-hydrolase domain enzymes (ABHDs). **(B)** LC3-I undergoes lipidation to form LC3-II, located on both sides of the phagophore membrane. LC3-II then recruits SQSTM1, which can be bound to the ubiquitinated cargo destined for degradation. HTT acts as a scaffold to support the function of SQSTM1. We predict that the palmitoylation of SQSTM1 and HTT plays an important role in their delivery of cargo to the phagophore membrane. Next, the phagophore expands around the cargo and closes to form a sealed autophagosome, which then fuses with a lysosome to form an autophagolysosome. All the contents inside this vesicle are degraded by hydrolytic enzymes, and LC3-II bound to the outside of the double-membrane is released back to the cytosol. **(C)** In HD, there is a defect at the cargo-loading step leading to an accumulation of empty autophagosomes and mHTT aggregates. We predict that this cargo-loading failure is at least partially the result of a reduction in SQSTM1 and HTT palmitoylation. Figure created with Biorender.

HD is a neurodegenerative disease with devastating affects on cognition and motor function^11^. HD is an autosomal dominant, monogenic disease caused by a mutation in exon 1 of the huntingtin gene (*HTT*) that encodes a cytosine-adenine-guanine (CAG) repeat expansion (*i*.*e*., 36 to 39 CAG repeats for partial HD penetrance; 40 or more repeats for full HD penetrance) resulting in a polyglutamine expansion in the huntingtin protein (HTT)^12,13^. In healthy conditions, HTT is ubiquitous and involved in many cellular processes^14^, including autophagy, in which HTT functions as a scaffold, binding to SQSTM1 and ubiquitinated cargo to support the function of SQSTM1 in cargo-loading^2,15–17^. However, cargo-loading is impaired in HD^9,10,17^, which may be at least partly due to reduced palmitoylation of SQSTM1 and mHTT^18^. We have shown that SQSTM1 palmitoylation plays an important role in cargo-loading and is required for SQSTM1 localization to autophagosomes^7,18,19^.

Palmitoylation, the most common form of S-acylation, is the reversible post-translational addition of a 16-carbon fatty acid to a cysteine residue via a thioester bond^20^ (Figure 1A). Due to its reversibility and hydrophobicity, palmitoylation allows selective and dynamic targeting of soluble proteins to membranes and thus plays a key role in regulating protein trafficking^20^. Palmitoylation is mediated by 23 palmitoyl acyltransferases (ZDHHCs) in humans, whereas the reverse reaction is mediated by acyl protein thioesterase (APT), palmitoylprotein thioesterase (PPT), and α/β-hydrolase domain (ABHD) enzymes^21^.

Notably, SQSTM1 palmitoylation is significantly reduced in the brains of HD patients and YAC128 mice^18,19^. In turn, increasing SQSTM1 palmitoylation using broad S-acylation inhibitor Palmostatin B improved HTT and SQSTM1 localization to autophagosomes^18,19^. Moreover, the Hayden group has shown that mHTT palmitoylation is reduced in HD mouse models and further decreases with increasing age and CAG repeat length^22,23^. HTT is palmitoylated by ZDHHC13 and ZDHHC17, and depalmitoylated by APT1 and APT2^22–25^. Importantly, HD is linked to decreased ZDHHC17 activity and increased APT1 activity^25,26^. We showed that Palmostatin B restores mHTT palmitoylation and thus mHTT aggregation and cytotoxicity in primary neurons derived from an HD mouse model^23^.

Like HTT, the Yang group has shown that SQSTM1 is depalmitoylated by APT1^18,27^. In turn, the Saudou group demonstrated that specific inhibition of APT1 using ML348 is protective in the R6/2 HD mouse model overexpressing N-terminal HTT lacking any HTT palmitoylation sites^26^. This suggests that increasing palmitoylation overall is protective. However, targeting palmitoylation in an HD mouse model expressing full length HTT may be more effective due to the relevance of HTT palmitoylation in HD and autophagy. Nevertheless, this study highlights that APT1 may be a therapeutic target in HD independent of HTT. Therefore, the identification of a small molecule which can increase SQSTM1 palmitoylation holds therapeutic promise for HD, rescuing cargo-loading in autophagy, and in turn, reducing mHTT aggregation and toxicity. Through a high-throughput screening of FDA-approved drugs, we identified Vorinostat, also known as suberoylanilide hydroxamic acid (SAHA), as a potential candidate for increasing SQSTM1 palmitoylation^28^. Vorinostat is primarily known as a blood-brain barrier (BBB) permeable inhibitor of lysine deacetylases (KDACs), more commonly known as histone deacetylases (HDACs), which catalyze the removal of acetyl groups (*i*.*e*., deacetylation) from lysine residues on proteins, predominantly histones^29,30^. Herein, we report a novel mechanism for Vorinostat, which significantly increases SQSTM1 palmitoylation leading to enhanced autophagic flux. Importantly, Vorinostat crosses the blood-brain barrier and restores SQSTM1 palmitoylation in the brains of YAC128 HD mice.

## 2. Materials and Methods

### 2.1 Cell Culture

HeLa cells were grown at 37°C in 5% CO_2_ in Dulbecco’s Modified Eagle Medium (DMEM; Wisent #319-015 CL) supplemented with 10% fetal bovine serum (FBS), 2 mM L-glutamine, 1 mM sodium pyruvate, 1X penicillin-streptomycin. Cells were seeded at 170,000 cells per 6 cm plate for biochemistry experiments and 75,000 cells per well onto 0.01 mg/mL poly-D-lysine coated coverslips (#1.5; VWR) in 6-well plates for microscopy experiments. The next day, cells were transfected with 10 μg of plasmid DNA using calcium phosphate precipitation and incubated for 2 hours. The transfection mix contained 1X HEPES-buffered saline (HBS) and 122 mM CaCl_2_. The media was replaced 2 hours post transfection. Where indicated, drugs were added to transfected cells immediately following the media change and cells were processed 18 hours post transfection (*i*.*e*., 16-hour drug incubation). For non-transfected cells, indicated drugs were added for 16 hours prior to processing. Drugs were added at the listed concentrations: 10 μM Vorinostat (Sigma #SML0061), 10 μM ML348 (Cayman #18523), 10 μM Palmostatin B (PalmB; Millipore #178501), 50 μM 2-bromopalmitate (2BP; Sigma #238422), and/or 100 nM Bafilomycin A1 (BafA1; Millipore #196000). All drugs were dissolved in dimethyl sulfoxide (DMSO), except for 2BP, which was dissolved in ethanol.

### 2.2 Click Chemistry Assay

Cell labeling and click chemistry were performed as previously described^31^. Briefly, 15-hexadecynoic acid (15-HDYA) (Click Chemistry Tools #CCT-1165) was saponified in potassium hydroxide and conjugated to fatty-acid free bovine serum albumin (BSA). Cells were then deprived of lipids in DMEM supplemented with 5% charcoal-stripped FBS (Wisent #080-710), 2 mM L-glutamine, 1 mM sodium pyruvate, 1X penicillin-streptomycin, and incubated with 15-HDYA for 4 hours, after which cells were lysed in modified RIPA buffer (50 mM HEPES pH 7.4, 150 mM NaCl, 2 mM MgCl_2_, 1% Igepal CA-630, 0.5% sodium deoxycholate, 0.1% SDS). Iodoacetamide was added to each sample to block free cysteine residues. Finally, lysates were subjected to click chemistry with biotin-azide (Click Chemistry Tools #CCT-1488), tris-hydroxypropyl triazolyl methylamine (THPTA; Click Chemistry Tools #CCT-1010), copper sulfate, and sodium ascorbate for 1 hour. The reaction was then ended by acetone precipitation. Since we are looking at total palmitoylated protein, the affinity purification step was skipped, and the protein was denatured at 95°C for 5 minutes in 1X sample loading buffer (SLB) with 20 mM dithiothreitol and stored at -20°C.

### 2.3 Acyl Biotin Exchange (ABE) Assay

The acyl biotin exchange (ABE) assay was performed as previously described^32^. Briefly, ground cortical brain tissue (approximately 30 mg) was sonicated (3×6 seconds at 20%) in ABE homogenization buffer (10 mM 10X phosphate buffer pH 7.4, 0.32 M sucrose, 1 mM EDTA, 6 M urea, 2.5% SDS) with 1X PIC and 20 mM methyl methanthiosulfate (MMTS). Alternatively, cells were sonicated (2×6 seconds at 20%) in lysis buffer (50 mM HEPES pH 7.4, 1 mM EDTA, 2% SDS) with 1X PIC and 20 mM MMTS. Lysates were incubated for 30 minutes at 50°C to block free cysteine residues with MMTS. Excess MMTS was removed by acetone precipitation and resuspended in 4SB (50 mM Tris pH 7.5, 5 mM EDTA, 4% SDS). Samples were then incubated with 0.7 M hydroxylamine and 1 mM HPDP-Biotin (APExBIO #A8008) for 1 hour rotating at room temperature to cleave thioester bonds and label newly freed cysteine residues with biotin. The reaction was ended by acetone precipitation and resuspended in lysis buffer without MMTS. Following protein quantification, 0.5-1 mg of protein was incubated in 150 mM NaCl dilution buffer (50 mM HEPES pH 7.0, 1% TX-100, 1 mM EDTA, 1 mM EGTA) with High-Capacity NeutrAvidin Agarose beads (Thermo #29202) for 3 hours rotating at 4°C to bind biotinylated proteins. Following two washes with 0.5 M NaCl dilution buffer and one wash with dilution buffer without NaCl, the protein was eluted from beads with elution buffer (dilution buffer, 250 mM NaCl, 0.2% SDS, 1% β-mercaptoethanol) over a 10-minute incubation at 37°C, then denatured at 95°C for 5 minutes in 1X SLB with 1% β-mercaptoethanol and stored at -20°C.

### 2.4 SDS-PAGE and Western Blotting

12% Bis-Tris polyacrylamide gels were run in 1X XT MOPS buffer (Bio-Rad #1610788) and transferred onto nitrocellulose (to detect SQSTM1) or a polyvinylidene fluoride (to detect LC3-II) membranes via semi-dry transfer using transfer buffer (192 mM glycine, 25 mM tris base, 20% methanol, 0.05% SDS) and the Transblot Turbo Transfer System. Primary antibodies rabbit anti-SQSTM1 (1:1000; Cell Signaling #D1Q5S) or rabbit anti-SQSTM1 (1:1000; Invitrogen #PA5-20839), rabbit anti-LC3A/B (1:1000; Cell Signaling #D3U4C), and mouse anti-VCP (1:1000; Proteintech #60316-1-Ig) were used to detect SQSTM1, LC3-II, and VCP, respectively. Secondary antibodies goat anti-rabbit Starbright 520 (1:2500; Bio-Rad #12005870), donkey anti-rabbit Alexa Fluor 790 (1:2500; Jackson ImmunoResearch #711-655-152), and goat anti-mouse Alexa Fluor 488 (1:2500; Jackson ImmunoResearch #115-545-003) were used. Tubulin-rhodamine (1:5000; Bio-Rad #12004166) was used to detect tubulin as the loading control. Alexa Fluor 680 Streptavidin (1:5000; Jackson ImmunoResearch #016-620-084) was used to detect biotinylated proteins.

### 2.5 Fluorescence Microscopy

For fixed cell imaging, cells were treated with 50 nM LysoTracker Deep Red (Invitrogen #L12492) for 1 hour prior to fixation. Cells were washed twice with 1X phosphate buffer saline (PBS), then fixed in 4% paraformaldehyde for 20 minutes at room temperature. Again, cells were washed twice with 1X PBS, then permeabilized in 1X PBS containing 0.1% TX-100, 1 μM CaCl_2_, 1 μM MgCl_2_ for 1 minute. Again, cells were washed twice with 1X PBS, then blocked in 0.2% gelatin for 30 minutes. Next, cells were incubated with the primary antibody for 1 hour. Rabbit anti-SQSTM1 (1:500; Invitrogen #PA5-20839) and mouse anti-HTT (1:500; Millipore #MAB2166) were used to detect SQSTM1 and HTT, respectively. Cells were washed twice with 0.2% gelatin, then the last step was repeated with the secondary antibody. Secondary antibodies goat anti-rabbit Alexa Fluor 488 (1:500; Invitrogen) and donkey anti-mouse Alexa Fluor 555 (1:500; Invitrogen) were used. Again, cells were washed twice with 0.2% gelatin, then incubated in 1 μg/mL DAPI (Sigma #D9542) for 20 minutes. Finally, cells were washed twice with 0.2% gelatin. The coverslips were mounted onto microscope slides (Fisher) using ProLong Gold Antifade Mountant (Invitrogen #P36934) and sealed with nail polish. Using a Nikon AXR laser scanning confocal microscope and its NIS-Elements software, Z-stack images were captured at 60X magnification. Next, 3D deconvolution was applied. Protein colocalization was determined by measuring Pearson Correlation Coefficient (PCC) between channels. For HeLa cells, PCC was determined from regions of interest manually drawn around single cells. For neurons, PCC was determined from regions of interest using the automated General Analysis 3 (GA3).

### 2.6 Molecular Docking

Molecular docking prediction was performed using the AutoDock Vina program^33^. Briefly, the protein PDBQT file was opened in AutoDockTools software to define the grid box dimensions. Molecular docking was then performed using AutoDock Vina via the command prompt, specifying the file path to the program and a folder containing the protein PDBQT file, the ligand PDBQT file, and a text file listing the PDBQT file names, grid box dimension values, energy range, and exhaustiveness. Energy range was set to 4 and exhaustiveness to 64. Running this program produced an “output.pdbqt” file containing the top ligand binding conformations, along with a “log.txt” file listing the corresponding binding affinities.

### 2.7 qPCR

Total RNA from cells was extracted using RNAZol RT (Bioshop #RNN190). Then, 500 ng of total RNA was used to prepare cDNA using High-Capacity cDNA Reverse Transcription Kit (Applied Biosystems #4368813). Next, cDNA was diluted 1:5, and qPCR was performed using primers and the PowerTrack SYBR Green Master Mix (Applied Biosystems #A46109). Each sample was run in triplicate. qPCR cycling conditions were 2 minutes at 95°C, followed by 40 cycles of 5 seconds at 95°C and 30 seconds at 60°C, then finally 15 seconds at 95°C, 1 minute at 60°C, and 15 seconds at 95°C. The forward and reverse primer sequences were synthesized by IDT and are listed in Table 1. *ACTB* and *GAPDH* were used as housekeeping genes. Primers for the ZDHHCs were previously published^34^. Primers for *SQSTM1, HTT, VCP, MAP1LC3B*, the histone deacetylases, the serine hydrolases, and the housekeeping genes were designed for this study.

**Table 1.**
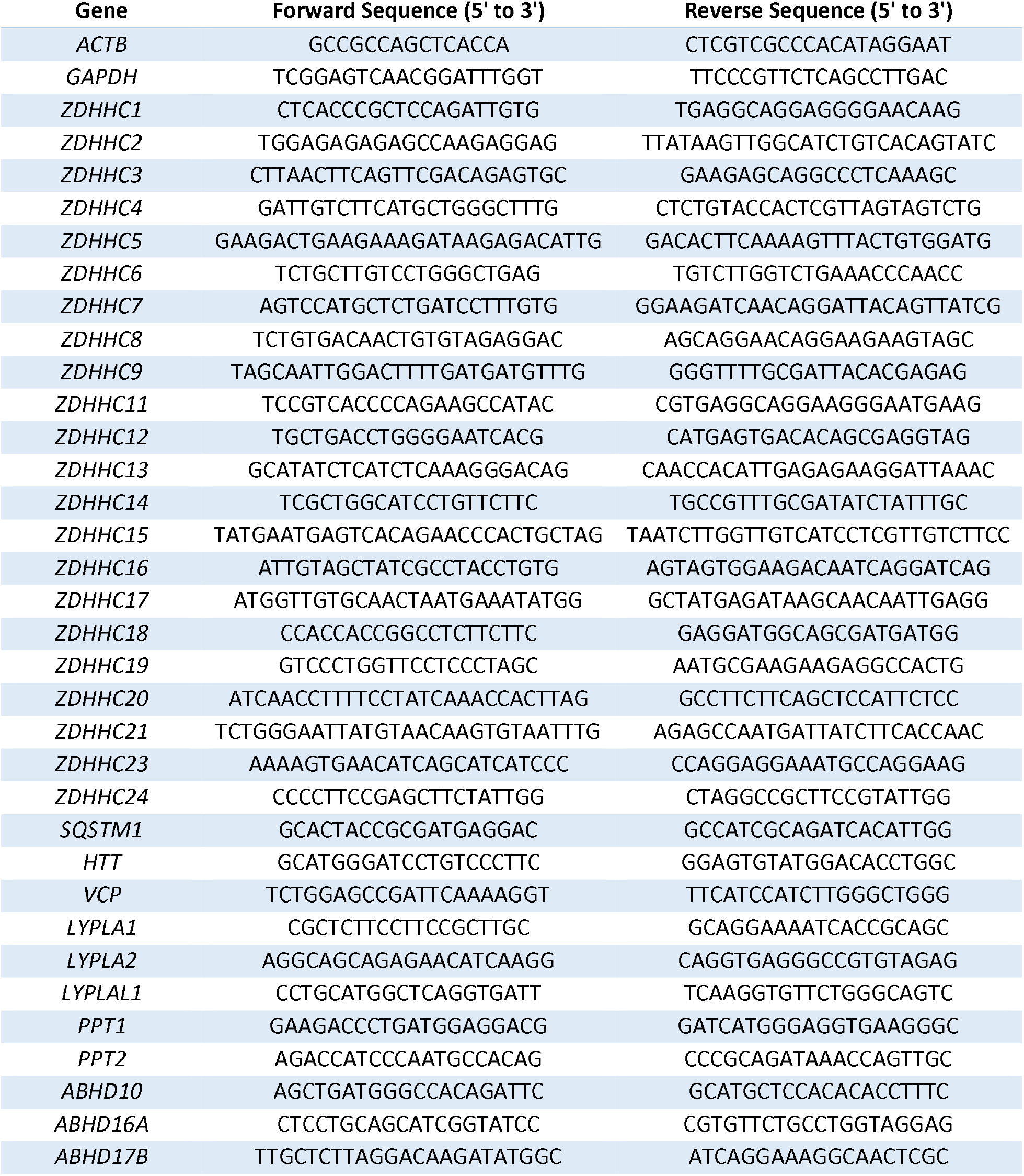
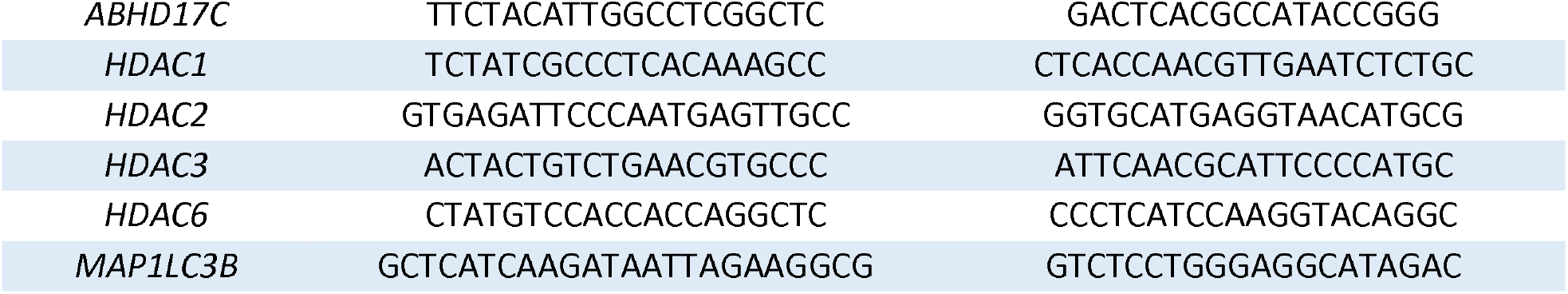
List of Human Primers for qPCR.

### 2.8 Mice

YAC128 transgenic mice on an FVB/N genetic background were obtained from in-house breeding colonies and were originally purchased from the Jackson Laboratory (Strain ID: 004938). Mice were housed in standard laboratory cages under a 12-hour light/dark cycle at 23 ± 1°C, with *ad libitum* access to food and water, and toys for enrichment. All mouse experiments were conducted in accordance with Animal Use Protocol #44651, approved by the UW Animal Care Committee. Mice were administered Vorinostat once every second day between 10 am and 12 pm, at a dose of 50 mg/kg through intraperitoneal (IP) injection (alternating right and left side) for four consecutive weeks for a total of 15 treatments^35^. Vorinostat was dissolved in DMSO at a concentration of 40 mg/mL and stored at -20°C. On the day of IP injection, the stock was diluted in corn oil to yield a working solution containing 18.75% DMSO and 81.25% corn oil, with a final Vorinostat concentration of 7.5 µg/µL. Control injections contained the same DMSO-to-corn oil ratio. Solutions were warmed to 37°C for 5 minutes and vortexed for 15 seconds prior to IP injection.

### 2.9 Mouse-Derived Primary Cortical Neurons

Primary cortical neurons were derived from YAC128 mouse embryos at embryonic day 17. Cortices were dissociated using papain and seeded on 1 mg/mL poly-L-lysine coated wells or glass coverslips (#1.5; Warner) in 6-well plates at 280,000 and 180,000 neurons per well for biochemistry and microscopy experiments, respectively. Neurons were plated in plating media [neurobasal media (Gibco #21103049) supplemented with 2.5% FBS, 2.5% horse serum, 2 mM L-glutamine, 1X penicillin-streptomycin, 2% B27]. The next day, plating media was replaced with neuronal media (neurobasal media supplemented 12 with 2 mM L-glutamine, 1X penicillin-streptomycin, 2% B27). Neurons were processed at 17 days *in vitro*.

### 2.10 Statistics

Statistical analyses and biological repeats are indicated in figure legends. Significant outliers identified using the Grubbs’ test were removed and are indicated in figure legends. GraphPad Prism 10 was used for all statistical analyses and graph preparation. Graphs are presented as the standard error of the mean.

## 3. Results

### 3.1 Vorinostat increases SQSTM1 palmitoylation but does not affect total protein palmitoylation

We identified Vorinostat as a regulator of palmitoylation through a high-throughput screening of FDA-approved drugs. To determine whether Vorinostat regulates autophagic receptor palmitoylation, we examined SQSTM1. Because SQSTM1 is highly dependent on proper localization for its function^6^, palmitoylation is a key mechanism regulating protein localization^20^, and SQSTM1 palmitoylation is decreased in HD^18,19^, we investigated whether Vorinostat could increase SQSTM1 palmitoylation. In turn, an ABE assay was performed to assess SQSTM1 palmitoylation. HeLa cells overexpressing SQSTM1-HA were treated with 10 μM Vorinostat, 10 μM PalmB, or vehicle. PalmB is a broad inhibitor of depalmitoylating enzymes^26^, whereas 2BP is a non-specific inhibitor of palmitoylation^36^. The absence of signal in the hydroxylamine (HAM-) lane confirms that without thioester bond cleavage, biotin-HPDP cannot be incorporated on free cysteine residues. Vorinostat treatment led to a significant increase in SQSTM1 palmitoylation (*p<0.05) (Figure 2A,B), while PalmB treatment led to no changes.

**Figure 2.**
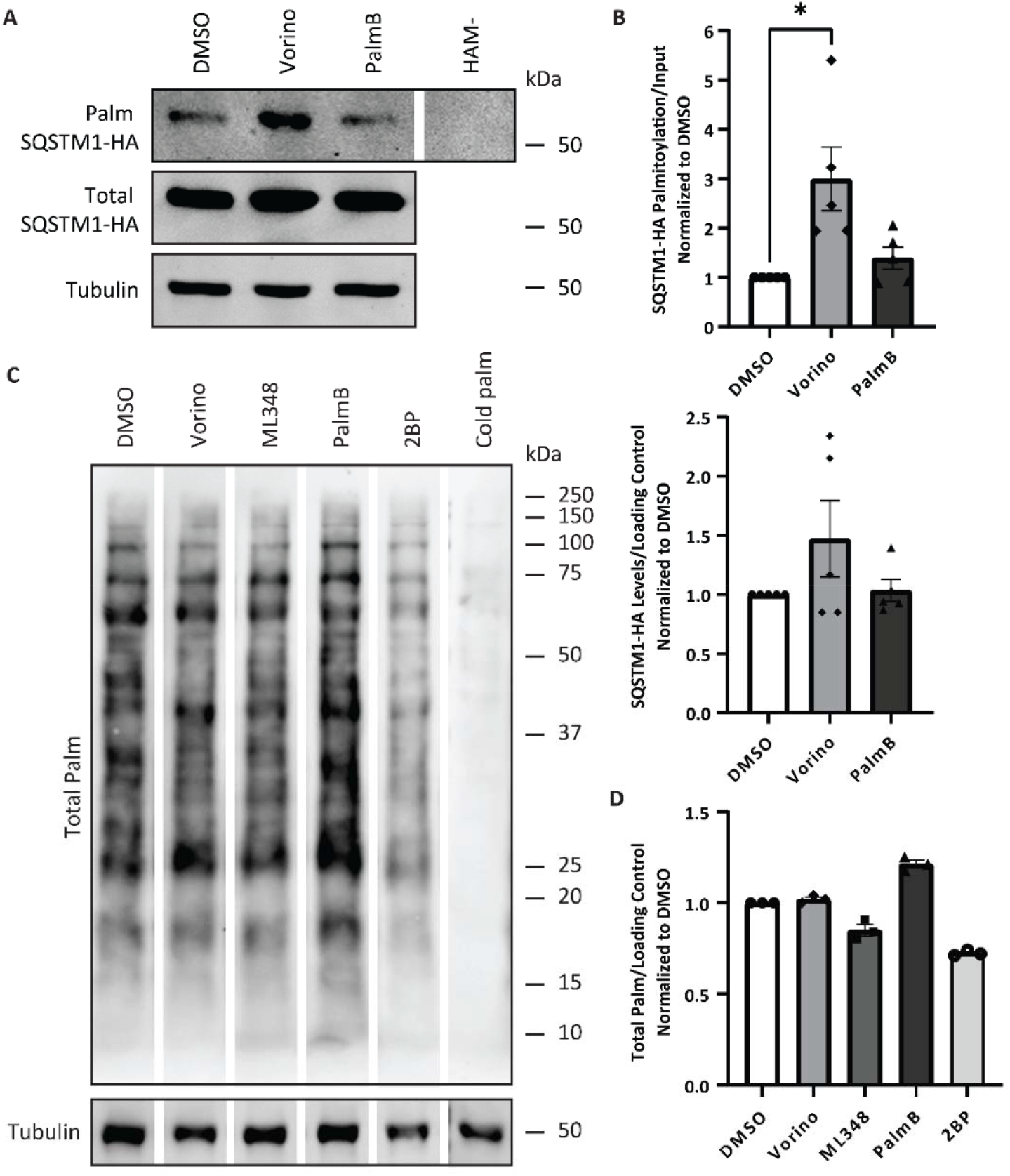
Vorinostat increases SQSTM1 palmitoylation but does not affect total protein palmitoylation. **(A)** SQSTM1 palmitoylation was detected via acyl biotin exchange assay using HeLa cells overexpressing SQSTM1-HA and treated with 10 μM Vorinostat, 10 μM PalmB, or vehicle. Composite of the same gel. **(B)** Quantification of (top) SQSTM1 palmitoylation and (bottom) SQSTM1 levels from five biological replicates. Kruskal-Wallis test followed by Dunn’s post-hoc analysis. **(C)** Total palmitoylated protein was detected via click chemistry using HeLa cells treated with 10 μM Vorinostat, 10 μM ML348, 10 μM PalmB, 50 μM 2BP, or vehicle in presence of 15-HDYA. Composite of the same gel. **(D)** Quantification of total palmitoylated protein from three biological replicates. Kruskal-Wallis test followed by Dunn’s post-hoc analysis.

Next, to compare total dynamic palmitoylation in HeLa cells treated with 10 μM Vorinostat, 10 μM ML348, 10 μM PalmB, 50 μM 2BP, or vehicle, we performed click chemistry assays using 15-HDYA. ML348 is a specific inhibitor for APT1^26^, the enzyme that depalmitoylates SQSTM1^18,27^. The absence of a signal in the cold palmitate (palm) lane confirms that without an alkyne group, biotin-azide cannot be added to the palmitate. Vorinostat treatment led to no changes in total protein palmitoylation (Figure 2C,D). ML348 and 2BP treatments led to a slight decrease in total protein palmitoylation, whereas PalmB led to a slight increase. Overall, these finding suggest that Vorinostat may promote palmitoylation of a subset of proteins, including SQSTM1.

### 3.2 Vorinostat crosses the blood-brain barrier and increases SQSTM1 palmitoylation in YAC128 HD mice

*In vitro*, Vorinostat increased SQSTM1 palmitoylation (Section 3.1). We next sought to determine whether Vorinostat crosses the blood-brain barrier and exerts the same effect *in vivo* in the YAC128 HD mouse model. We also examined valosin-containing protein (VCP). VCP has been identified in palmitoyl-proteomic studies^37,38^ and plays important roles in autophagy^39^. Moreover, VCP is the major pathogenic protein in multisystem proteinopathy (MSP)^40^ and one of the several pathogenic proteins implicated in amyotrophic lateral sclerosis (ALS)^41^. Therefore, we investigated whether Vorinostat also increases VCP palmitoylation in addition to SQSTM1. 22-month-old YAC128 mice were administered 50 mg/kg Vorinostat or vehicle via intraperitoneal injection once every second day for four consecutive weeks. Subsequently, an ABE assay was performed using the mouse cortices. Vorinostat treatment in YAC128 mice led to a significant increase in SQSTM1 and VCP palmitoylation (*p<0.05) compared to YAC128 vehicle-treated and WT Vorinostat-treated mice (Figure 3A,B). Vorinostat treatment in WT mice did not affect SQSTM1 and VCP palmitoylation, nor total protein palmitoylation in both WT and YAC128 mice.

**Figure 3.**
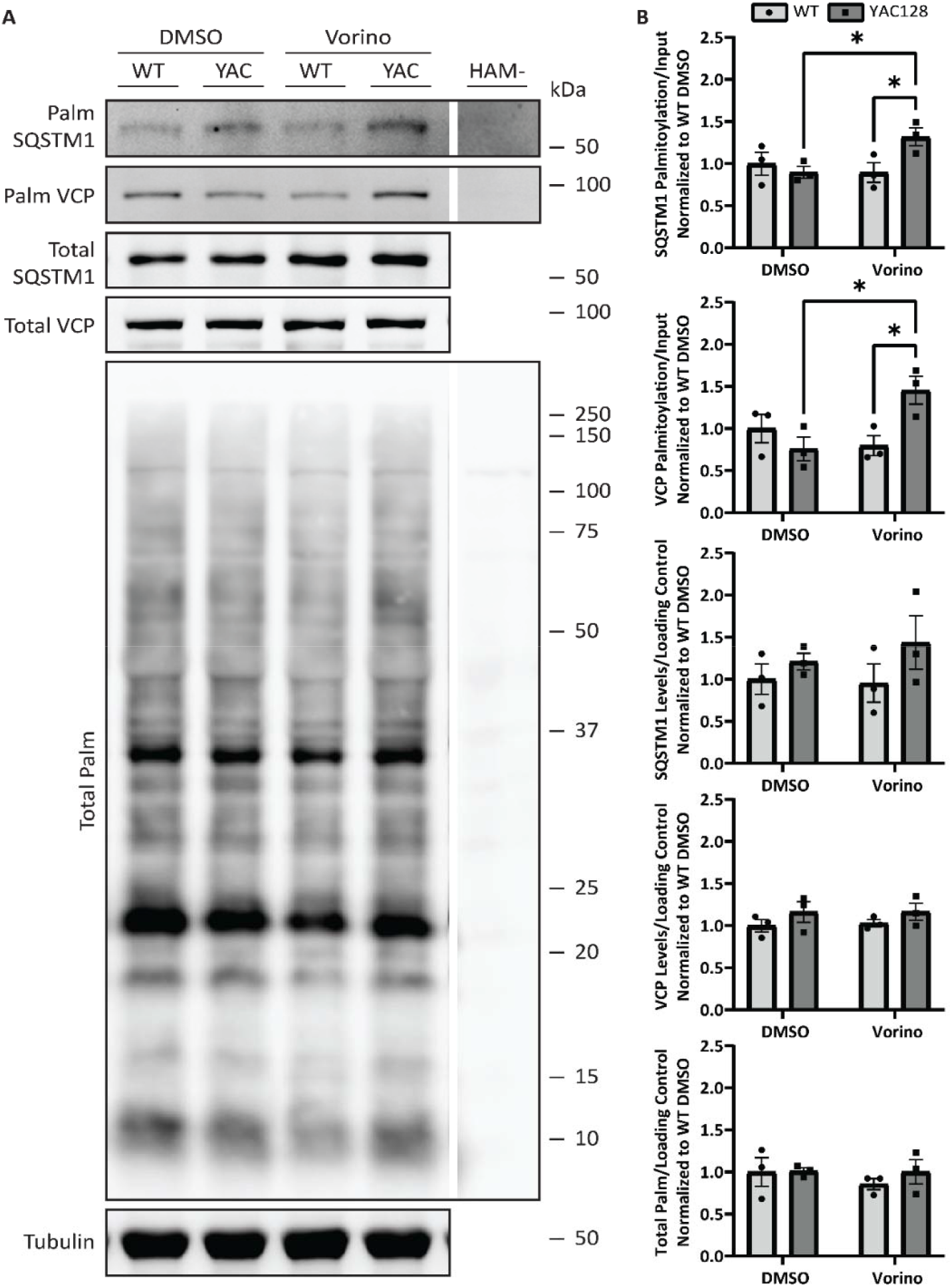
Vorinostat crosses the blood-brain barrier and increases SQSTM1 palmitoylation in YAC128 mice. **(A)** SQSTM1 palmitoylation was detected via acyl biotin exchange assay using cortices from YAC128 mice that were administered 50 mg/kg Vorinostat or vehicle every other day for 4 consecutive weeks. Composite of the same gel. **(B)** Quantification of (top) SQSTM1 palmitoylation, (second) VCP palmitoylation, (third) SQSTM1 levels, (fourth) VCP levels, and (bottom) total palmitoylated protein from three biological replicates. Two-way ANOVA test followed by Fisher’s LSD post-hoc analysis (*p<0.05).

### 3.3 Vorinostat induces autophagy and redirects HTT to lysosomes

Given that Vorinostat increases SQSTM1 palmitoylation in YAC128 mice, we next examined its effect on autophagy. In autophagy, SQSTM1 is degraded in autophagolysosomes, and while some of LC3-II located on the inner surface is also degraded, the majority located on the outer surface is recycled back to LC3-I, the inactive delipidated form of LC3^42^. This makes SQSTM1 and LC3-II excellent markers of autophagy, such that increased autophagic flux is indicated by reduced SQSTM1 levels and elevated LC3-II levels^42^. BafA1 is a late-stage autophagy inhibitor that prevents autophagosome-lysosome fusion leading to an increase in both SQSTM1 and LC3-II levels^42^. BafA1 is used to confirm that the effect of the drug treatment on autophagic flux is indeed a result of increased autophagy^42^. HeLa cells were treated with 10 μM Vorinostat, 10 μM ML348, 10 μM PalmB, or vehicle, with or without BafA1. Vorinostat treatment led to a significant decrease in SQSTM1 levels (*p<0.05) (Figure 4A,B), with a concomitant significant increase in LC3-II levels (*p<0.05), indicating autophagy has been induced and SQSTM1 is being degraded in the autophagolysosome. ML348 and PalmB treatment led to no changes in SQSTM1 or LC3-II levels. All BafA1 treatment conditions led to a significant increase in LC3-II (**p<0.01) and SQSTM1 (***p<0.001).

**Figure 4.**
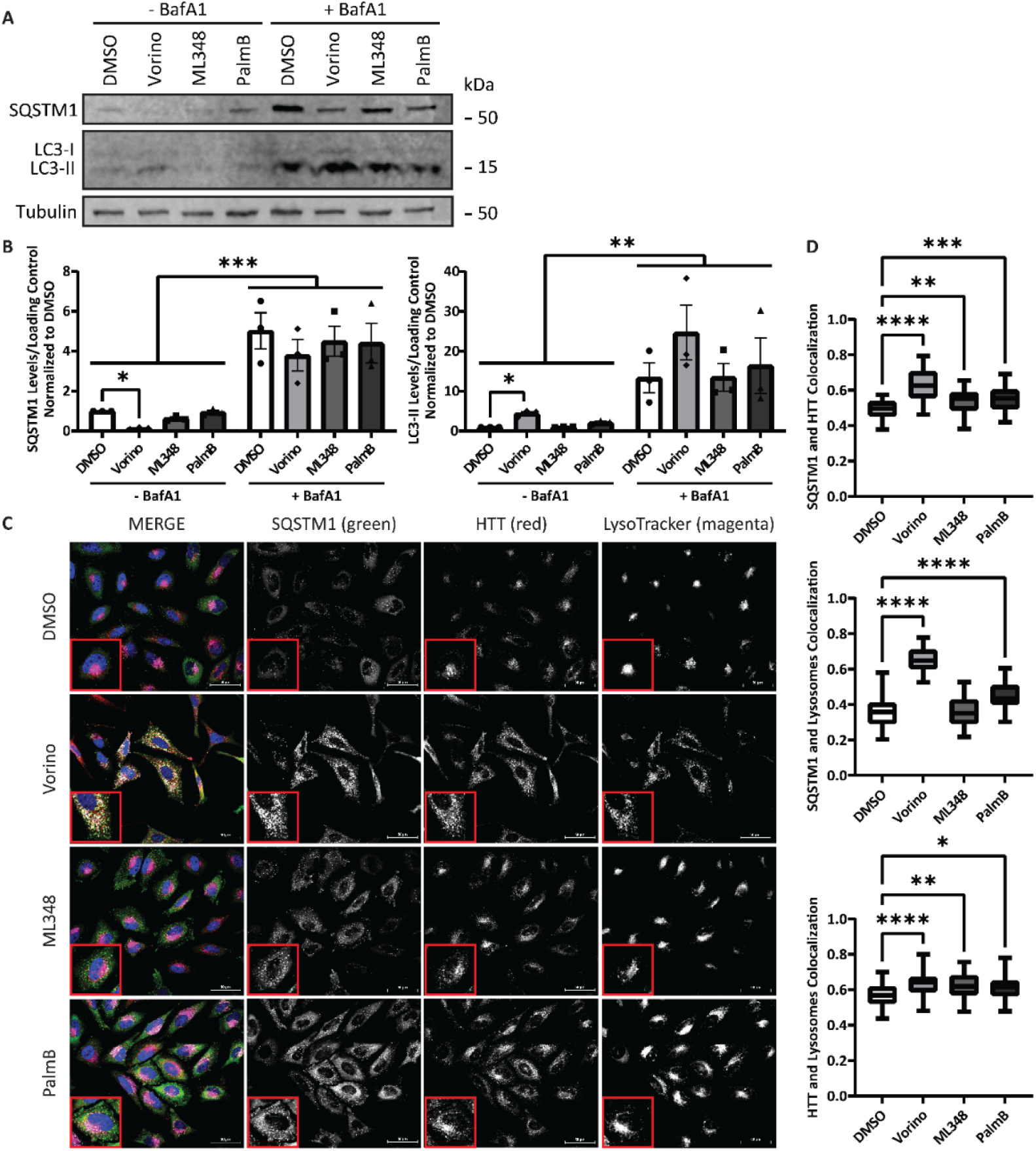
Vorinostat increases autophagic flux. **(A)** SQSTM1 and LC3-II levels were detected in HeLa cells treated with 10 μM Vorinostat, 10 μM ML348, 10 μM PalmB, or vehicle, with or without BafA1. **(B)** Quantification of (left) SQSTM1 levels and (right) LC3-II levels from three biological replicates. Kruskal-Wallis test followed by Dunn’s post-hoc analysis (*p<0.05) within +/-BafA1 conditions. Two-way ANOVA test followed by Fisher’s LSD post-hoc analysis (**p<0.01, ***p<0.001) across BafA1 conditions. **(C)** SQSTM1, HTT, and lysosomes were visualized via confocal immunofluorescence microscopy using fixed HeLa cells treated with 10 μM Vorinostat, 10 μM ML348, 10 μM PalmB, or vehicle. **(D)** Quantification of (left) PCC between SQSTM1 and HTT, (middle) PCC between SQSTM1 and lysosome, and (right) PCC between HTT and lysosome, 15 cells per condition from three biological replicates. One-way ANOVA test followed by Dunnett’s post-hoc analysis (*p<0.05, **p<0.01, ***p<0.001, ****p<0.0001). Box and whiskers (min to max).

Furthermore, an indicator of functional autophagy is the colocalization of SQSTM1, HTT, and lysosomes, which suggests that SQSTM1 is successfully delivering cargo (*e*.*g*., mHTT) with the support of HTT to autophagosomes, which fuse with lysosomes to form autophagolysosomes. Therefore, increased colocalization between these proteins and organelles, as measured by the PCC, indicates enhanced autophagic flux, as the presence of SQSTM1 and HTT in lysosomes signifies their successful delivery. Consequently, HeLa cells were treated with 10 μM Vorinostat, 10 μM ML348, 10 μM PalmB, or vehicle and visualized by fixed cell confocal immunofluorescence microscopy. When treated with Vorinostat, ML348, or PalmB, SQSTM1 and HTT colocalization significantly increased, with a more pronounced increase elicited by Vorinostat (****p<0.0001) (Figure 4C,D). When treated with Vorinostat or PalmB, SQSTM1 and lysosome colocalization significantly increased, with a more pronounced increase detected in Vorinostat treated cells (****p<0.0001). ML348 had no effect on SQSTM1 and lysosome colocalization. Finally, when treated with Vorinostat, ML348, or PalmB, HTT and lysosome colocalization significantly increased, with a greater increase induced by Vorinostat (****p<0.0001).

To confirm the effect in HD, primary cortical neurons derived from WT and YAC128 mice were treated with 10 μM Vorinostat, 10 μM ML348, 10 μM PalmB, or vehicle and visualized by fixed cell confocal immunofluorescence microscopy. In WT cortical neurons treated with Vorinostat, SQSTM1 and HTT colocalization was not affected (Figure 5A,C). However, in YAC128 cortical neurons treated with Vorinostat, SQSTM1 and HTT colocalization significantly increased (***p<0.001). Similarly, SQSTM1 and lysosome colocalization was not affected with Vorinostat treatment in WT cortical neurons but significantly increased (**p<0.01) in YAC128 cortical neurons. In addition, HTT and lysosome colocalization, which includes both HTT and mHTT in YAC128 cortical neurons, was not affected with Vorinostat treatment in WT cortical neurons but significantly increased (****p<0.0001) in YAC128 cortical neurons, which is especially relevant in this context where colocalization suggests that mHTT will be degraded in autophagolysosomes. When treated with ML348, WT cortical neurons showed significantly increased SQSTM1 and HTT colocalization (**p<0.01), but not HTT and lysosome or SQSTM1 and lysosome colocalization. Whereas YAC128 cortical neurons showed significantly increased SQSTM1 and HTT colocalization (***p<0.001), as well as HTT and lysosome colocalization (**p<0.01), but not SQSTM1 and lysosome colocalization. When treated with PalmB, WT cortical neurons had significantly increased SQSTM1 and HTT colocalization (**p<0.01), as well as HTT and lysosome colocalization and ****p<0.0001), but not SQSTM1 and lysosome colocalization. Whereas YAC128 cortical neurons showed significantly increased HTT and lysosome colocalization (***p<0.001), but not SQSTM1 and HTT or SQSTM1 and lysosome colocalization. Overall, these results suggest that increasing total palmitoylation in YAC128 cortical neurons directs HTT to SQSTM1 and lysosomes; however, only Vorinostat was effective at redirecting both SQSTM1 and HTT together and to lysosomes, and to a much greater extent.

**Figure 5.**
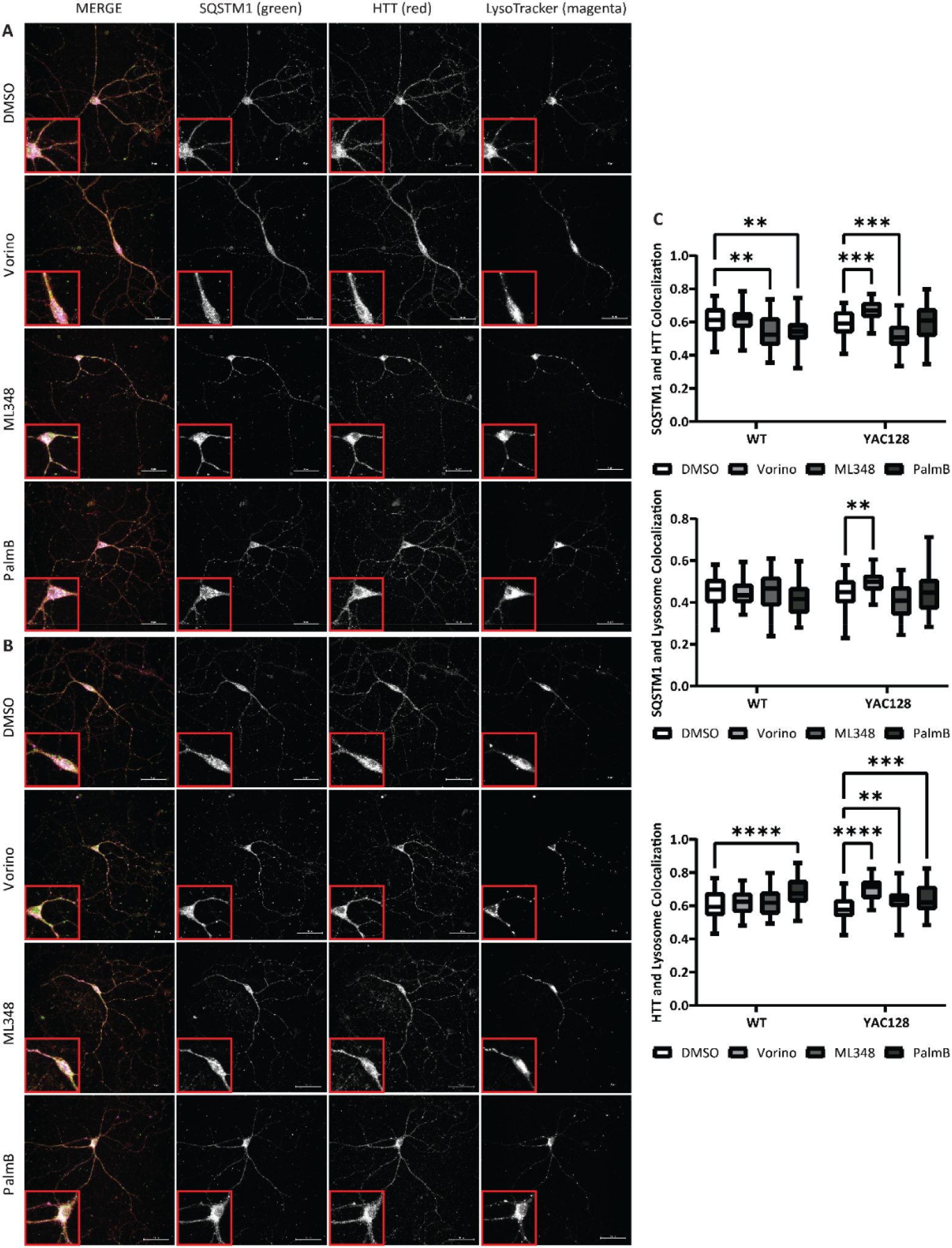
Vorinostat redirects HTT and SQSTM1 to lysosomes in HD neurons. SQSTM1, HTT, and lysosomes were visualized via confocal immunofluorescence microscopy using fixed primary cortical neurons derived from **(A)** WT and **(B)** YAC128 mouse embryos treated with 10 μM Vorinostat, 10 μM ML348, 10 μM PalmB, or vehicle. **(C)** Quantification of (top) PCC between SQSTM1 and HTT, (middle) PCC between SQSTM1 and lysosome, and (bottom) PCC between HTT and lysosome, on an average of 15 neurons per condition from three biological replicates. Two-way ANOVA test followed by Dunnett’s post-hoc analysis (**p<0.01, ***p<0.001, ****p<0.0001). Box and whiskers (min to max).

### 3.4 Vorinostat may function as a depalmitoylation inhibitor and transcriptional regulator

SQSTM1 has been shown to be depalmitoylated by APT1^18,27^. Thus, our initial theory was Vorinostat may function via inhibition of APT1, thereby increasing SQSTM1 palmitoylation. To further investigate, molecular docking experiments were performed using AutoDock Vina. These results indicate that ML348, an inhibitor specific to APT1^26^, and Vorinostat have a deep fit into the binding pocket of APT1, with ML348 exhibiting a stronger binding affinity (-10.6 kcal/mol) compared to Vorinostat (-8.0 kcal/mol) (Figure 6A). We also looked at APT2 for comparison and found that ML349, an inhibitor specific to APT2^26^, and Vorinostat have a deep fit into the binding pocket of APT2, with ML349 exhibiting a stronger binding affinity (-12.9 kcal/mol) compared to Vorinostat (-8.0 kcal/mol). For context, the more negative the value, the stronger the binding affinity.

**Figure 6.**
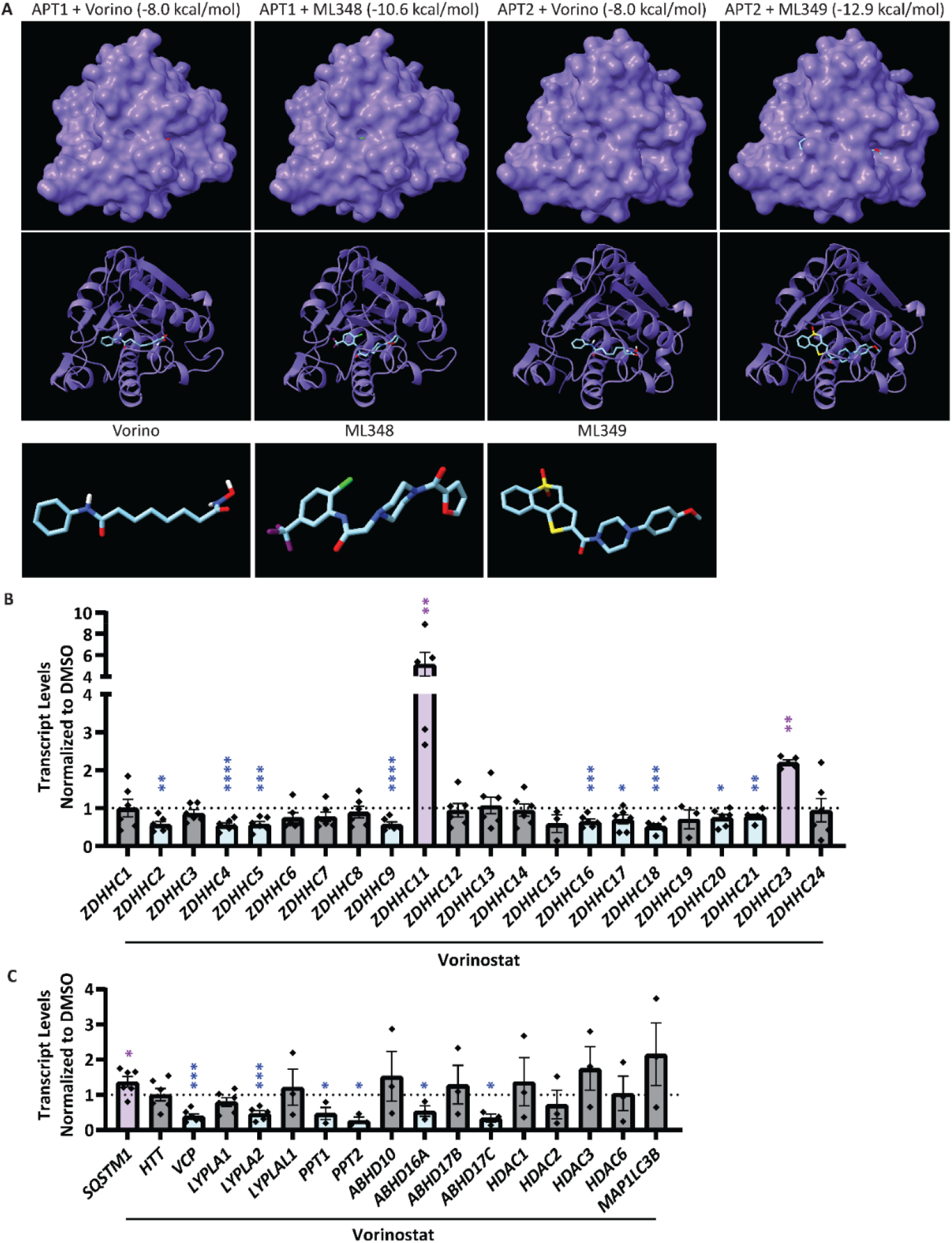
Vorinostat may function as a depalmitoylation inhibitor and transcriptional regulator. **(A)** Vorinostat and ML348 molecular docking and binding affinity on APT1 and APT2 were predicted using AutoDock Vina. **(B)** *ZDHHC1-24* transcript levels were detected via qPCR using HeLa cells treated with 10 μM Vorinostat or vehicle. *ZDHHC2,4,5,9,11,16,17,18,20,21,23* transcript levels are sensitive to Vorinostat treatment. One significant outlier (SO) removed from ZDHHC11 and ZDHHC23. Two-tailed unpaired *t*-test (*p<0.05, **p<0.01, ***p<0.001, ****p<0.0001). **(C)** *SQSTM1*; *HTT; VCP*; *LYPLA1,2*; *LYPLAL1*; *PPT1,2*; *ABHD10,16A,17A,17B,17C*; *HDAC1,2,3,6*; and *MAP1LC3A,B* transcript levels were detected via qPCR using HeLa cells treated with 10 μM Vorinostat or vehicle. *SQSTM1, VCP, LYPLA2, PPT1, PPT2, ABHD16A*, and *ABHD17C* transcript levels are sensitive to Vorinostat treatment. One SO removed from LYPLA2. Two-tailed unpaired *t*-test (*p<0.05, ***p<0.001).

Alternatively, Vorinostat is already known as an HDAC inhibitor^29,30^, so its function could be through increasing the transcription of palmitoylating enzymes or decreasing the transcription of depalmitoylating enzymes. To further investigate, qPCR was performed using HeLa cells treated with 10 μM Vorinostat or vehicle to quantify the relative transcript levels of *ZDHHC1-24, SQSTM1, HTT, VCP, LYPLA1* (APT1), *LYPLA2* (APT2), *PPTs, ABHD*s, *HDAC*s, and *MAP1LC3*s (Figure 6B,C). Compared to vehicle, treatment with Vorinostat led to significantly increased *ZDHHC11, ZDHHC23*, and *SQSTM1* transcript levels, as well as significantly decreased transcript levels of *ZDHHC2, ZDHHC4, ZDHHC5, ZDHHC9, ZDHHC16, ZDHHC17, ZDHHC18, ZDHHC20, ZDHHC21, VCP, LYPLA2, PPT1, PPT2, ABHD16A*, and *ABHD17C*. Notably, *HTT, LYPLA1*, and *ZDHHC19* transcript levels were unaffected by Vorinostat treatment. ZDHHC22 had amplification in the Vorinostat condition, but not vehicle condition, whereas ABHD17A and MAP1LC3A showed no amplification in either condition.

## 4. Discussion

Autophagy is disrupted in HD at various stages^8^. In particular, a failure at the cargo-loading step has been linked to SQSTM1’s inability to deliver cargo to the phagophore membrane for degradation resulting in an accumulation of empty autophagosomes^9,10^. Since palmitoylation plays an important role in protein localization to membranes^20^, we predicted that the autophagic dysfunction seen in HD is linked to reduced SQSTM1 palmitoylation detected in the brains of HD patients and YAC128 mice^18,43^. Consequently, we sought to promote mHTT intracellular degradation, thereby preventing cytotoxicity and neuronal death, by restoring SQSTM1 palmitoylation. Through a high-throughput screening of FDA-approved drugs, we identified blood-brain permeable Vorinostat as a candidate to increase SQSTM1 palmitoylation.

We found that Vorinostat treatment significantly increases SQSTM1 palmitoylation but does not affect total protein palmitoylation both *in vitro* (Figure 2) and *in vivo* (Figure 3). Since the biochemical assays for our *in vivo* work were conducted on cortical tissue, this confirms that Vorinostat successfully crosses the blood-brain barrier. Notably, PalmB did not increase SQSTM1 palmitoylation *in vitro*. This may be because PalmB is a broad inhibitor of depalmitoylating enzymes^26^. Since many palmitoylating enzymes undergo regulatory palmitoylation (*e*.*g*., ZDHHC6 is palmitoylated by ZDHHC16^44^ and ZDHHC5 is palmitoylated by ZDHHC20^45^), broad inhibition of depalmitoylating enzymes could disrupt the function of palmitoylating enzymes, potentially explaining the lack of effect of PalmB on SQSTM1 palmitoylation.

With an increase in SQSTM1 palmitoylation induced by Vorinostat treatment, we then observed a corresponding increase of autophagic flux. We found that Vorinostat treatment significantly decreases SQSTM1 levels and increases LC3-II levels (Figure 4). SQSTM1 and LC3-II are autophagy markers, and their respective decrease and increase in levels suggests increased autophagic flux, since SQSTM1 is degraded in autophagolysosomes, whereas the majority of LC3-II is recycled back into the inactive delipidated form of LC3^42^. Increased autophagic flux was confirmed with Vorinostat plus BafA1 treatment, a late-stage autophagy inhibitor^42^, which significantly increased SQSTM1 and LC3-II levels.

Furthermore, with an increase in SQSTM1 palmitoylation induced by Vorinostat treatment, we then observed a corresponding enhancement of autophagy. We found that Vorinostat treatment significantly increased SQSTM1, HTT, and lysosome colocalization in HeLa cells (Figure 4), as well as primary cortical neurons derived from YAC128 mice, but not WT mice (Figure 5). In HeLa cells, this suggests enhanced SQSTM1 and HTT delivery to autophagosomes. In YAC128 primary cortical neurons, increased SQSTM1, HTT, and lysosome colocalization suggests improved delivery of HTT to autophagosomes for subsequent degradation in autophagolysosomes. Notably, the observation that Vorinostat had an effect in YAC128 but not in WT primary cortical neurons is important as it suggests that Vorinostat may exert its effects selectively under pathological conditions, reducing the risk of potential off-target effects. The effects of ML348 and PalmB on SQSTM1, HTT, and lysosome colocalization were neither as strong, consistent, nor specific as those of Vorinostat, suggesting that Vorinostat may be a more effective drug in HD.

Initially, we predicted that Vorinostat increases SQSTM1 palmitoylation by acting as a depalmitoylation inhibitor, specifically of APT1, the depalmitoylating enzyme of SQSTM1. Consequently, we aimed to better understand Vorinostat’s ability to inhibit depalmitoylating enzymes through molecular docking experiments to quantify Vorinostat’s binding affinity to APT1 and APT2 (Figure 6). Although Vorinostat had a deep fit into the binding pockets of both APT1 and APT2, its binding affinity to either was weaker than that of the specific inhibitors, ML348 and ML349. However, prior experiments indicated that Vorinostat was more effective than ML348 at increasing SQSTM1 palmitoylation, thus prompting us to reassess our previous thinking. Vorinostat is also a known HDAC inhibitor, promoting histone acetylation or transcription factor acetylation, thereby increasing transcription^29,30^. Either way, Vorinostat could increase the transcription of *ZDHHC* genes, leading to increased SQSTM1 palmitoylation through an alternative pathway. Hence, Vorinostat may be acting via a dual mechanism, as summarized in Figure 7.

**Figure 7.**
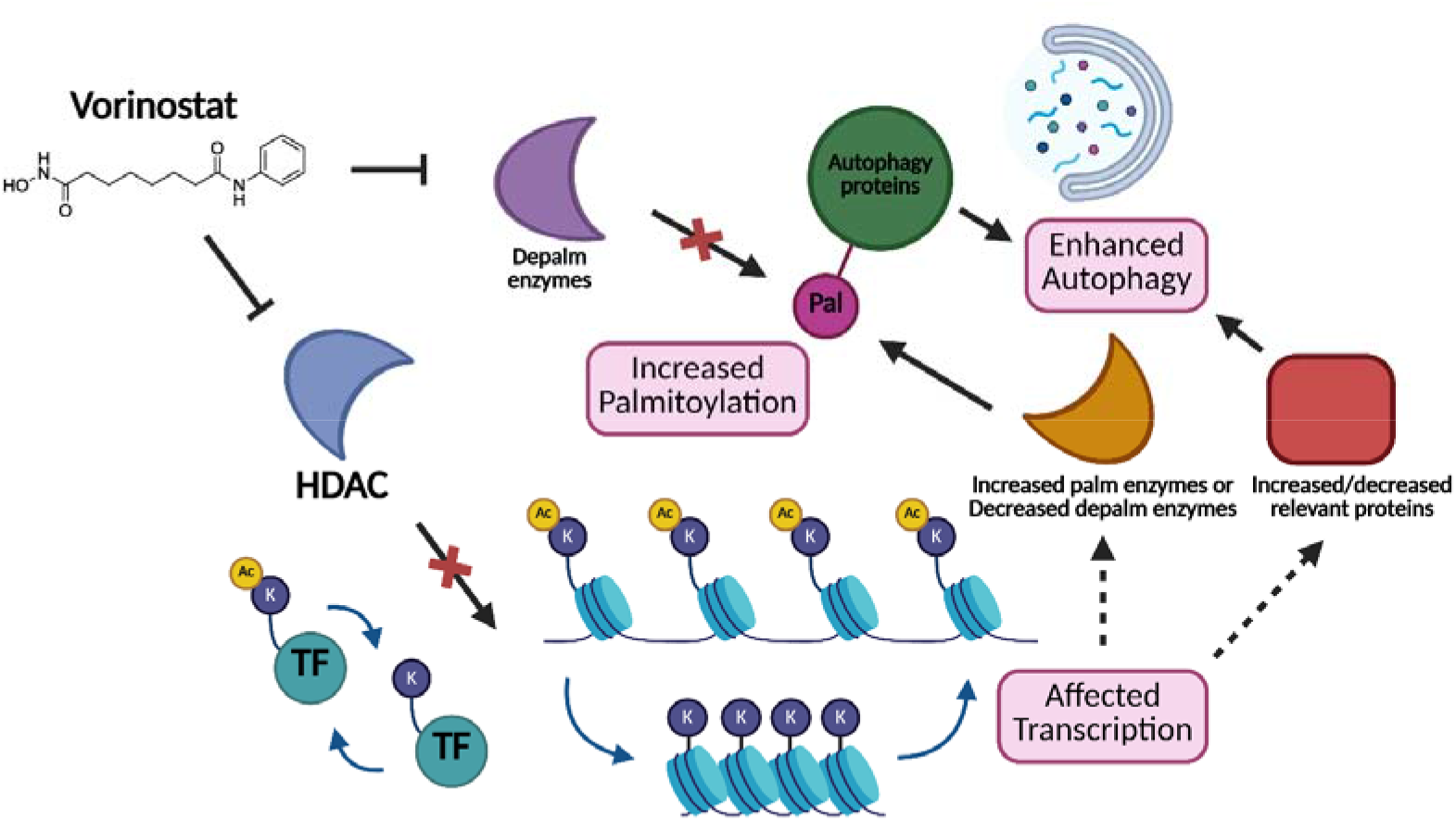
Predicted Vorinostat mechanism. Vorinostat may be working through two concurrent mechanisms to increase autophagy protein palmitoylation: inhibition of depalmitoylating (depalm) enzymes and inhibition of acetylation enzymes (HDAC). Inhibition of depalmitoylating enzymes prevents the depalmitoylation of SQSTM1, thus increasing SQSTM1 palmitoylation. In contrast, inhibition of HDAC prevents deacetylation of histones, thereby increasing their acetylation, promoting active transcription by opening chromatin. Alternatively, inhibition of HDAC prevents deacetylation of transcription factors (TF), thus increasing their acetylation, also promoting active transcription (K represents a lysine residue protruding from the histone, and Ac represents an acetyl group). This may increase transcription of palmitoylating (palm) enzymes or decrease the transcription of depalmitoylating enzymes, overall increasing autophagy protein palmitoylation. Alternatively, this may increase or decrease transcription of relevant proteins, such as SQSTM1, which may also enhance autophagy. Figure generated on BioRender.

The next step was to confirm whether Vorinostat increases transcription of relevant proteins by quantifying transcript levels via qPCR. We found that Vorinostat significantly increases *SQSTM1* transcript levels compared to vehicle, suggesting that Vorinostat may promote autophagy. However, Vorinostat plus BafA1 did not lead to an increase in SQSTM1 levels (Figure 4). It could be that Vorinostat plus BafA1 leads to the accumulation of detergent-insoluble SQSTM1, as described by the Yang group^27^, which may limit detection of SQSTM1 via western blot. In addition, we found that Vorinostat significantly increases *ZDHHC11* and *ZDHHC23* transcript levels compared to vehicle. Work by the Liu group suggests that ATG2A is palmitoylated by ZDHHC11 and its *depalmitoylation* is required for proper autophagosome formation^46^. In contrast, studies by the Kim group suggest that MCOLN3/TRPML3 is palmitoylated by ZDHHC11 and its *palmitoylation* is required for proper autophagosome formation^47^. Minimal connections between ZDHHC23 and autophagy have been established. Therefore, it remains unclear how Vorinostat-induced increases in the *ZDHHC11* and *ZDHHC23* transcript levels might have a role in the increase and enhanced autophagy we observed following Vorinostat treatment. Furthermore, we found that Vorinostat decreases *LYPLA2* transcript levels which encode APT2. APT2 depalmitoylates HTT, and HTT palmitoylation is reduced in HD mouse models^23^. Therefore, reducing APT2 may increase HTT palmitoylation, thereby lowering mHTT aggregation and cytotoxicity. Although increased transcript levels do not necessarily equate to increased protein levels, they serve as a good representation of potential regulatory changes at the transcriptional level and can provide insight into upstream effects of Vorinostat treatment.

### 4.1 Vorinostat and Huntington disease

Vorinostat has been tested in HD mouse models previously, but in the context of its HDAC inhibiting properties and fixing transcriptional dysregulation found in HD^35,48^. The Bates group treated the R6/2 HD mouse model with Vorinostat and found it restored motor impairment, reduced mHTT aggregates in the cortex, and restored *Bdnf* transcript levels^35,48^. Similarly, the Saudou group found restored BDNF trafficking with APT1 inhibition by ML348 in the R6/2 model^26^. BDNF is implicated in HD pathogenesis^49^. In R6/2 mice, *Bdnf* transcript levels decline during disease progression^50^; accordingly, BDNF over-expression was found to improve HD phenotypes^51^. The Bates group also found that Vorinostat decreases HDAC2 and HDAC4 levels *in vivo*^*48*^. In R6/2 mice, the group reported that HDAC4 associates with mHTT in a polyQ-dependent manner and colocalizes with cytoplasmic inclusions, and knockdown of HDAC4 reduced cytoplasmic aggregate formation, restored *Bdnf* transcript levels, rescued membrane properties of MSNs and of corticostriatal synaptic transmission, improved motor and neurological impairment, and extended the lifespan of R6/2 mice^52^. Of note, the R6/2 model only expresses the very N-terminus of HTT, which does not include any palmitoylation sites^24^ or SQSTM1 binding sites^17^.

To further understand the mechanism of HDAC4, the Cristea group studied its interaction in the context of HD through multiomics and proposed a potential model of the role for HDAC4 in HD^53^. One notable finding from their study is the interaction between HDAC4 and VCP, which they also found has minimal interaction in the Q20 WT mice and maximal interaction in the Q140 HD mice, an interaction which increases with age. Combining this with our findings on VCP, it is possible that the increased interaction between HDAC and VCP, and likely VCP acetylation^54^, interferes with VCP palmitoylation, thereby disrupting its normal function. By deceasing HDAC4 levels with Vorinostat, VCP function may be restored.

### 4.2 VCP and Huntington disease

VCP is involved in autophagy initiation^39^ and HD^55^. The Qi group identified VCP as a mHTT-binding protein on mitochondria, where it drives excessing mitophagy, or the autophagic degradation of mitochondria^55^. Because palmitoylation is a key regulator of protein localization^20^, restoring VCP palmitoylation could potentially correct its mislocalization, reducing VCP accumulation on mitochondria in HD mouse models. Thus, we looked at whether Vorinostat also rescues VCP palmitoylation in addition to SQSTM1 palmitoylation and found that, *in vivo*, Vorinostat treatment significantly increases VCP palmitoylation.

### 4.3 Future Directions

Although studies have demonstrated positive outcomes of Vorinostat treatment in HD mouse models, the broader consequences of HDAC inhibition remain an important consideration. Determining the specific mechanism by which Vorinostat mediates these positive effects should be a priority, as this knowledge could guide the design of more targeted small molecules. However, it is also possible that Vorinostat’s multi-functional nature contributes to its broad impact on various HD phenotypes, whether through restoring protein palmitoylation, correcting transcriptional dysregulation, or both. In this study, we reported that Vorinostat crossed the blood-brain barrier and increased SQSTM1 palmitoylation *in vivo*, as well as enhanced autophagy *in vitro*, facilitating HTT loading into autophagolysosomes for degradation.

## Author Contributions

YA and DDOM conceived of experiments. FA completed preliminary work (not included). YA completed the experiments in Figures 2-6. FA helped complete the SQSTM1 palmitoylation experiment in Figure 2. MR helped complete the qPCR experiments in Figure 6. AD helped complete the docking experiments in Figure 6. FR helped with animal studies including preparation of primary neuronal cultures. YA prepared the first draft. YA prepared Figures 1-7. YA and DDOM edited the final draft.

## Acknowledgements

We would like to acknowledge that the University of Waterloo resides on the traditional territory of five Indigenous communities, including the Ho-de-no-sau-nee-ga (Haudenosaunee), Mississaugas of the Credit First Nation, Anishinabewaki, Attiwonderonk (Neutral), and Mississauga peoples, which is situated on the Haldimand Tract, a land granted to the Six Nations, encompassing six miles on either side of the Grand River. This land is home to many past, present, and future Indigenous peoples.

